# Vocal recognition of partners by female prairie voles

**DOI:** 10.1101/2024.07.24.604991

**Authors:** Megan R. Warren, Larry J. Young, Robert C. Liu

## Abstract

Recognizing conspecifics is vitally important for differentiating kin, mates, offspring and social threats.^1^ Although often reliant upon chemical or visual cues, individual recognition across the animal kingdom is also facilitated by unique acoustic signatures in vocalizations.^2–4^ However, amongst the large *Muroidea* superfamily of rodents that encompasses laboratory species amenable to neurobiological studies, there is scant behavioral evidence for individual vocal recognition despite individual acoustic variation.^5–10^ Playback studies have found evidence for coarse communicative functions like mate attraction and territorial defense, but limited finer ability to discriminate known individuals’ vocalizations.^11–17^ Such a capacity would be adaptive for species that form lifelong pair bonds requiring partner identification across timescales, distances and sensory modalities, so to improve the chance of finding individual vocal recognition in a *Muroid* rodent, we investigated vocal communication in the prairie vole (*Microtus ochrogaster*) – one of the few socially monogamous mammals.^18^ We found that the ultrasonic vocalizations of adult prairie voles can communicate individual identity. Even though the vocalizations of individual males change after cohabitating with a female to form a bond, acoustic variation across individuals is greater than within an individual so that vocalizations of different males in a common context are identifiable above chance. Critically, females behaviorally discriminate their partner’s vocalizations over a stranger’s, even if emitted to another stimulus female. These results establish the acoustic and behavioral foundation for individual vocal recognition in prairie voles, where neurobiological tools^19–22^ enable future studies revealing its causal neural mechanisms.

**Highlights:**

- Muroid rodents can display individual vocal recognition
- Adult prairie vole USVs are more variable across individuals than social experience
- Individual vole identity can be decoded from their vocalizations
- Carefully constructed protocol sustains vole’s interest in vocal playback
- Female prairie voles behaviorally recognize their mate’s vocalizations

## INTRODUCTION

Rodents are a neurobiologically accessible taxa that emit communicative vocalizations,^5,23^ which have been suggested by some to carry individual information,^6–8^ but see ^24^. However, empirical evidence for individual vocal recognition in laboratory rodents like rats and mice is lacking. One reason may be that in many laboratory rodents, social recognition memory is thought to be largely chemically mediated.^25,26^ Moreover, many argue that rodent USVs just convey a vocalizer’s arousal state,^27,28^ so playback studies^11–14^ have typically looked just for communicative function rather than testing for individual vocal recognition. Additionally, habituation to sound playback is commonly observed in those studies, which makes it difficult to infer any sustained differential preference for one individual’s vocalizations over another’s – perhaps explaining why discrimination studies have been relatively rare.^12,15^

One laboratory rodent model for which individual vocal recognition may have been highly beneficial is the prairie vole, which forms socially monogamous pair bonds between mated adults.^18^ These enduring bonds require partner recognition across many timescales and distances, likely involving multiple sensory modalities. Adult prairie voles do in fact emit vocalizations in both the audible and ultrasonic frequency range.^29,30^ We investigated the possibility that these vocalizations communicate individual identity, both acoustically and behaviorally, by designing a behavioral paradigm centered around the social experience to establish a lifelong pair bond.

## RESULTS

We focused on recording and testing ultrasonic vocalizations (USVs) from adult prairie voles. We allowed pairs of male and female voles to freely interact within a recording arena for 30 minutes (**Figure 1A**). Individual males (n=7) were first placed with an unfamiliar stimulus female (Day 0), and then a different unfamiliar female (Day 1) who was to become his mated partner. Following this ‘pre-cohabitation’ condition, the males were co-housed with their soon-to-be partners for seven days in the colony room to solidify their pair bond. Pairs were then separated for 24 hours and brought back for both a playback study (Day 9, see below) and a ‘post-cohabitation’ social interaction (Day 9). During each encounter, we recorded video and audio, extracting 141,788 total vocal segments across 21 recordings (6752±3618 segments/recording, mean±standard deviation, unless otherwise noted).

**Figure 1:**
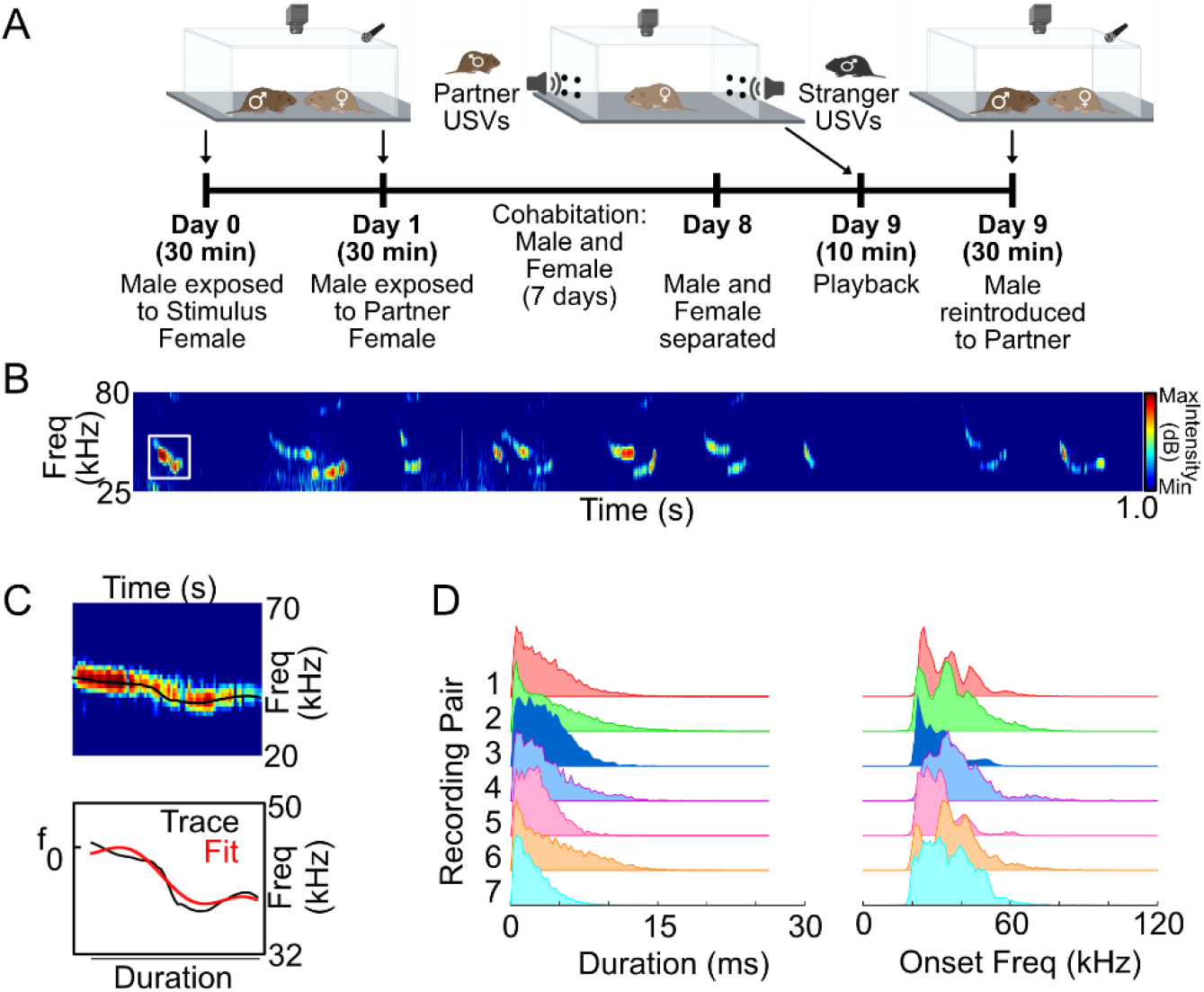
Experimental protocol. **A)** Timeline of experiments, created with BioRender.com. **B)** Spectrogram of 1 second of vocal activity from a Day 1 recording. **C)** Schematic of vocal feature extraction. Top shows vocalization spectrogram, with warmer colors indicating louder sound. Black line traces the actual fundamental frequency, which is reproduced in the Bottom panel along with the frequency fitted (red) according to a sinFM model. **D)** Distributions of duration **(Left)** and onset frequency **(Right)** parameters from the sinFM fits for all vocal segments from each animal pair recorded on Day 1.

We fit vocal segments to a combination of linear and sinusoidal frequency modulated (sinFM) tones whose parameters (**Figure 1B**) are known to modulate auditory responses in rodents.^31^ The distributions of each of the fitted parameters (**Figure 1C**) were coarsely similar, yet often distinct statistically, from individual to individual (Pairwise Kolmogorov-Smirnov tests, with Bonferroni correction. Duration: 19/21 significant comparisons; Onset frequency: 21/21 significant comparisons; uncorrected p<0.0024). To visualize acoustic differences between the vocal repertoires emitted in each individual session, we embedded each segment’s six-dimensional parameters into a two-dimensional vocal space (see Methods, **Figure 2A**).

**Figure 2:**
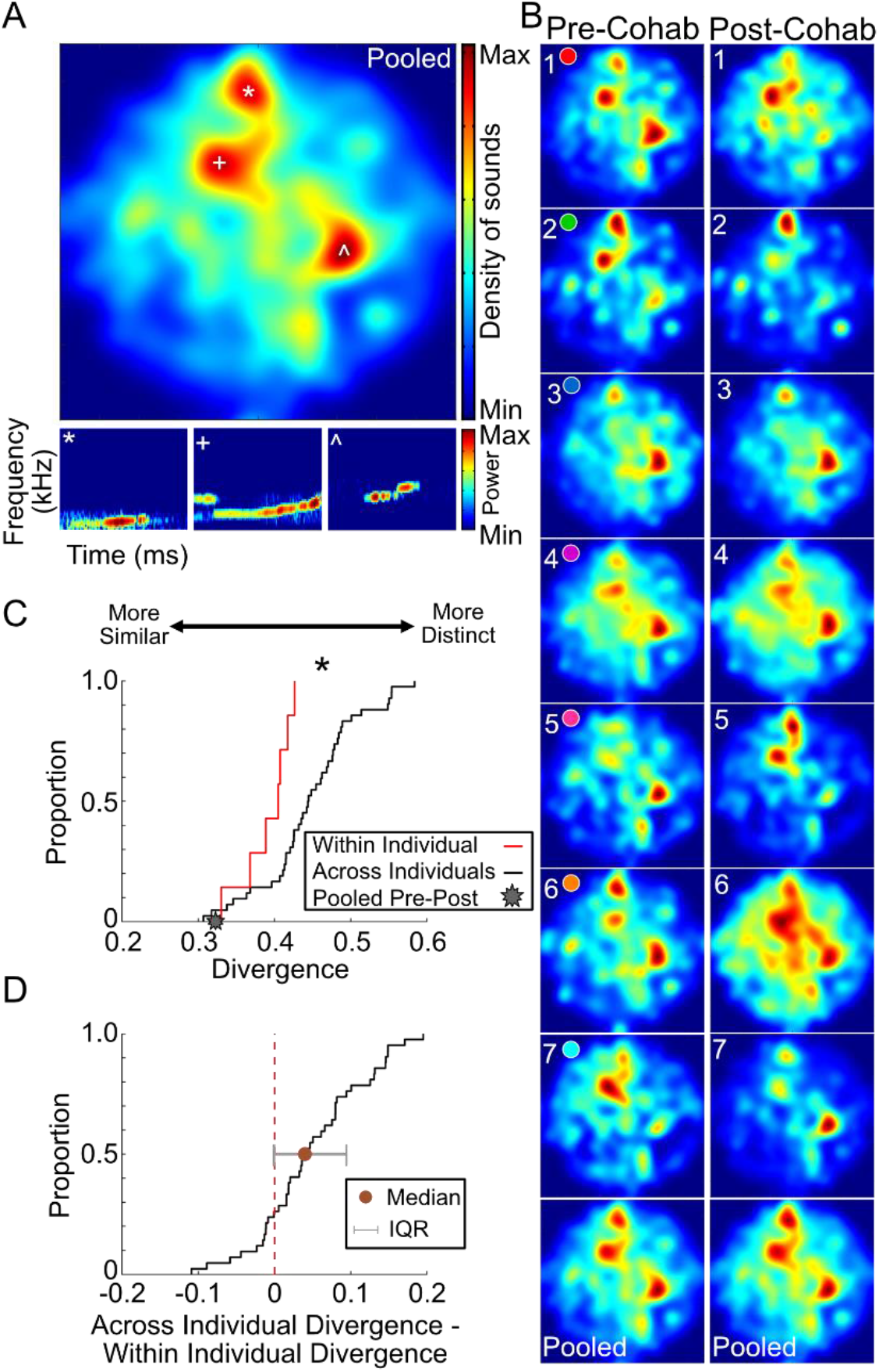
Vocalizations differ more between individuals than across experiences. **A) (Top)** 2D heat map of sinFM features across all emitted vocal segments after dimensionality reduction (t-SNE). **(Bottom)** Example vocalizations that fall into respective t-SNE peaks. **B)** Individual t-SNE maps for each of the seven unique male-female pairing **(Rows)** for pre-cohabitation **(Left column)** and post-cohabitation **(Right column)** recordings. Colored circles indicate different pairs (consistent across figures). **(Bottom row)** Pre- and post-cohabitation t-SNE maps pooled across all pairs. **C)** Cumulative distribution of Jensen-Shannon divergences computed for pairwise comparisons of the t-SNE maps within all seven individuals from pre-to-post cohabitation (red line, n=7), across all possible different individuals whether pre- or post-cohabitation (black line, n=7*6*2), and across pre-to-post cohabitation for vocal segments pooled over all individuals (star, n=1). **D)** Cumulative distribution of voles’ (n=7) across-individual Jensen-Shannon divergences (included in black line in C) normalized by its own within-individual divergence (included in red line in C). Median (brown dot with interquartile range in gray) of this distribution is significantly higher than 0, indicating individuals are more similar vocally to themselves than to others.

Session-specific vocal spaces were variable across cohabitation experience and across different individual pairs (columns and rows of **Figure 2B**, respectively). We quantified this variability using the Jensen-Shannon divergence.^32^ All possible pairwise comparisons between different individuals’ pre- and/or post-cohabitation repertoires ranged from 0.307 to 0.584 (0.447±0.063). Interestingly though, when vocal segments were pooled across all individuals from the same context, the overall spaces for pre- and post-cohabitation vocalizations were largely similar (**Figure 2B, bottom row**), with a divergence of 0.326 that was near the lowest end of all empirically measured comparisons across individuals (**Figure 2C, star**, z-test, z=1.93, p=0.03). Hence, even though individual male-female pairs vocalized differently depending on their cohabitation experience, which vocalizations they emitted in those contexts was, on average, not simply dictated by experience.

The lack of a discernible systematic effect of pair bonding on vocalizations could imply that each male-female encounter simply produces an independent collection of USVs, which are as variable across individual pairs as they are across cohabitation experience. If true, such randomness would not be conducive to individual vocal recognition. To test for this, we considered whether divergences derived from comparing the pre- and post-cohabitation vocal spaces *within the same* individuals were any different than those calculated from all possible comparisons *across different* individuals (**Figure 2C**). We found that these two distributions were significantly different (Kolmogorov-Smirnov test, KS=0.62, p=0.01), with the latter shifted to larger divergences than the former. In fact, the residual across-individual divergence for subjects once their own within-individual pre-to-post divergence was subtracted out was significantly different than zero (signedrank, z=3.86, p<0.001; **Figure 2D**) and positive. Hence, the vocalizations of different prairie voles tended to be more distinct from each other than those emitted by the same prairie voles between their initial and pair bonded social interactions.

Greater variability in the vocalizations of different male-female pairs could provide the basis for individuals to be recognized from their vocal emissions. We investigated this at the acoustic level by training a classifier (**Figure 3A**) to predict individual identity from vocalizations. Given the variability seen across contexts, we limited ourselves to the vocalizations emitted from the same social context. For post-cohabitation, our classifier successfully predicted the identity of individual males with an accuracy of 0.74±0.15, which was significantly above chance (t=128.7, p<0.001, chance=0.14, **Figure 3B)**. Hence, vocalizations from the same context were emitted with sufficiently individualized features. These were presumably dominated by the male’s emissions^33–35^ but to control for the fact that different females were present during each of the post-cohabitation recordings with the partner, we separately trained a classifier on the vocal segments from Day 0, when all males were exposed to the same stimulus female. We were still able to predict identity with an accuracy of 0.85±0.02, which was again well above chance (t=986.4, p<0.001, **Figure 3C**). Thus, any female vocal contributions were unlikely to provide the only identifying information in the recording. Male prairie voles must then emit vocalizations with enough acoustic individuality that the emitter can in principle be accurately recognized, regardless of who he vocalizes to.

**Figure 3:**
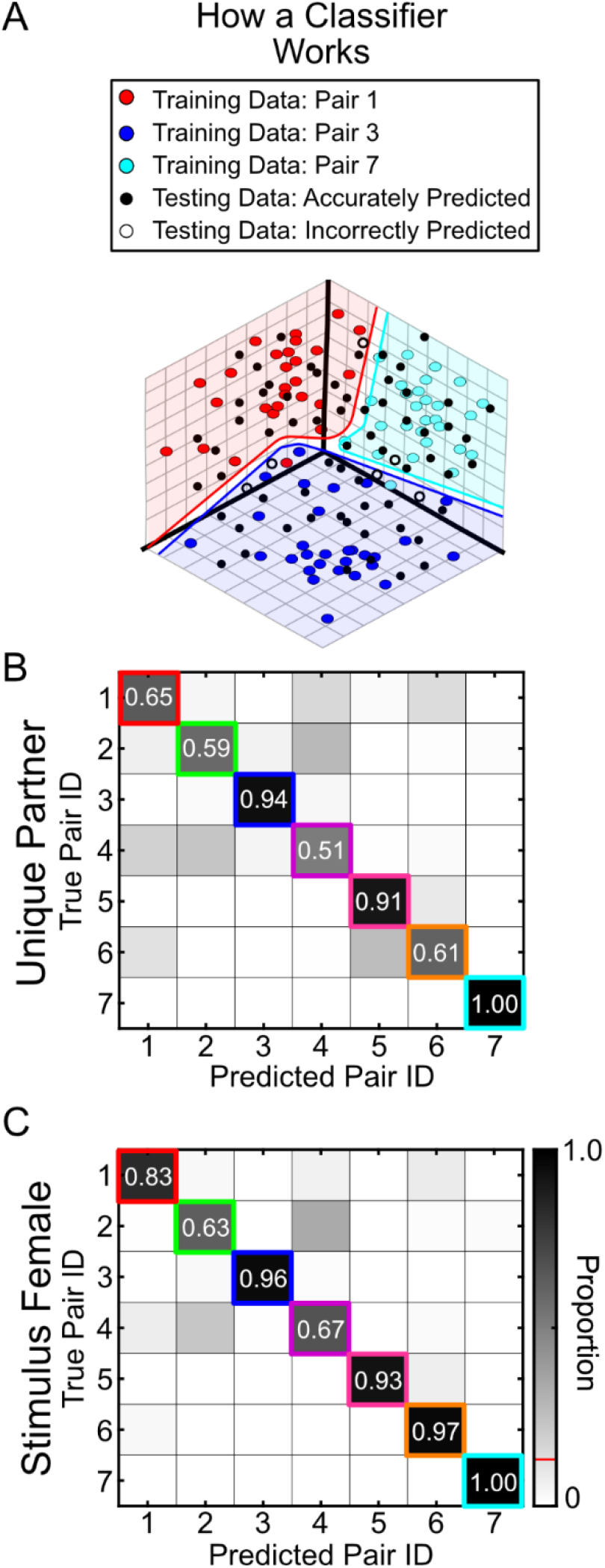
Male identity can be determined from sinFM Features. **A)** Schematic of how a single-vector multi-class classifier works to predict the class of unlabelled test data (black dots) after supervised training (red, blue and cyan dots). Colored regions depict corresponding hyperplanes. **B)** Classifier accuracy in determining which male emitted individual vocal segments based on their sinFM acoustic features while males are socializing to their to-be partners (Day 1). Boxes on the identity line show correct identity predictions. Values off the identity line show incorrect predictions. **C)** Same as **B)**, but vocal signals were from times when all males interacted with the same stimulus female (Day 0).

Finally, we wanted to know whether acoustic distinguishability in principle translates in practice to behavioral recognition of the vocalizing male by his female partner. We devised a playback study to present each female prairie vole with the Day 0 USVs emitted *to the stimulus female* by either their own partner or by a stranger male (who was partnered with a different female). We generated sound files (**Figure 4A**) in which every 30 seconds the stimulus emitted from a given speaker was either USVs from the appropriate male (partner or stranger, speaker side randomized and counter-balanced per playback session) or background noise (**Figure 4B**). Hence, there were times when only USVs from one male was playing, times when USVs from both males were playing, and times when no USVs were playing.

**Figure 4:**
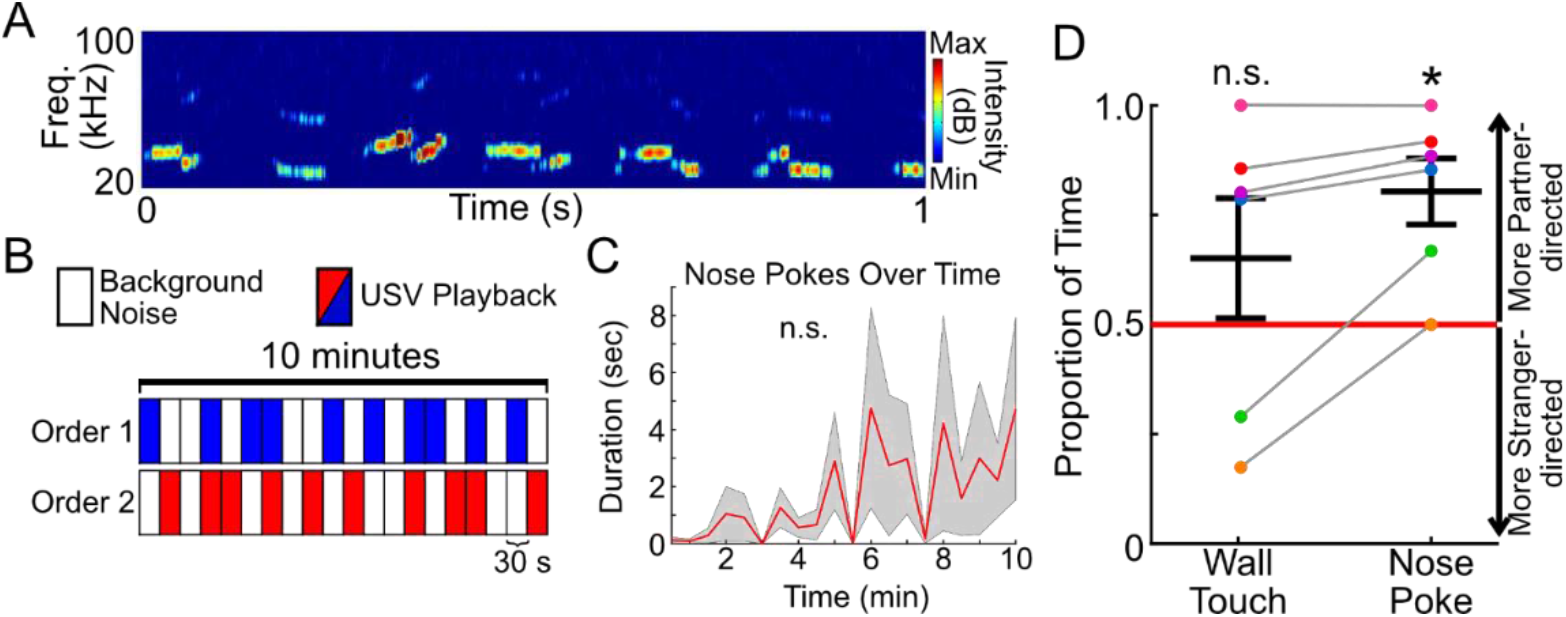
Female prairie voles show individual vocal recognition of their mates. **A)** Representative spectrogram showing 1 second of audio data recorded during free social interaction between a male and a female. **B)** Schematic of playback organization. During playback, sounds were concurrently played back from two speakers on opposite ends of the arena; one file containing USVs emitted by the experimental female’s partner, and one containing USVs from an unfamiliar male. Speaker side (left or right) and playback ordering (1 vs 2) were randomized. **C)** Total time spent nose-poking per 30 second playback interval. Red line is mean across individuals; gray is standard error. Change over time assessed with repeated-measures ANOVA. **D)** Proportion of time females spent touching the wall **(Left)** or nose poking **(Right)** towards the USVs of her partner during playback. 50% = chance level. Dots colored by recording ID. Thick black lines represent mean and standard error of the mean. T-tests compared distributions to chance level.

We blindly assessed two types of sound-directed behavior: time when the female was touching the wall where either speaker was positioned, and times when the female poked its nose through holes in the wall to get closer to the speaker. We found no significant change in the more active “nose poke” behavior over time (F=1.17, p=0.31), indicating a lack of habituation to sound playback (**Figure 4C**). Using “wall touches,” the partner-vs stranger-directed behavior was similar (t=-1.14, p=0.32; **Figure 4D**).

However, we found that females spent a significantly larger proportion of time nose poking towards the emission of her partner’s USVs than the emission of a stranger’s USVs (t=-6.44, p<0.003). Thus, even when playing back novel USVs emitted to another female, pair bonded females could consistently identify the vocal signals of their male partners, indicating that male prairie vole calls carry individualized signatures that are behaviorally relevant for the vocal recognition of partners.

## DISCUSSION

Rodents are not generally known for an ability to recognize individuals based on their vocalizations, yet by leveraging an ethologically grounded playback paradigm in prairie voles, we demonstrate that males emit acoustically distinctive USVs that females are able to use behaviorally to identify their partner. Our discovery brings monogamous rodents into the diverse collection of species in which individual vocal recognition has been found behaviorally,^2^ spanning amphibians to birds and mammals – but no *Muroid* rodents. Importantly, as a neurobiologically accessible rodent species,^19–22^ the prairie vole adds the opportunity to uncover the causal neural mechanisms by which the auditory processing of identifying vocal features elicits individual-directed behavioral responses.

Unique acoustic signatures in the vocalizations of common laboratory rodents have been debated previously,^7,8,10,24^ yet even if they exist, studies did not reveal whether animals use them to identify individuals. That gap stems in part from a lack of robust effects of a sender’s vocalizations on a receiver’s behavior. Prior playback studies showed that calls (often versus silence) can elicit initial investigation of the speaker, but behavioral responses habituate rapidly.^11,14^ Adult calls can be discriminated after operant training,^36^ but whether that occurs endogenously for individual identification is unknown. Wild female mice can distinguish non-kin vocalizations from those of kin they had not actually heard vocalize, suggesting a heritable group level rather than individual level recognition.^15^

Similarly, behavioral studies of pup vocalizations suggest that dams can at least discriminate calls from pups of their own litter^37^ or of different sexes,^38^ but neither necessarily implies identifying individuals. In fact, while prairie vole pups vocalize at higher rates than pups of similar species,^39,40^ their calls become more stereotyped between early infancy and later stages (e.g., P6 to P16),^41^ making it unclear whether adult vole vocalizations contain sufficient individuality. Our acoustic analysis and decoding results indicate they do.

Individual vocal recognition presumably evolved to facilitate differential responses to distant conspecifics with whom interactions would have divergent costs or benefits. For animals that form lifelong pair bonds, recognizing a mate’s vocalizations before they are seen and discriminating them from that of a stranger could mean the difference between welcoming home a co-parent or defending their offspring from an intruder. The adaptive benefit of multimodal partner recognition may be one reason why we were able to observe a robust sign of a female prairie vole’s high interest in the playback of her partner’s vocalizations in the absence of seeing or smelling him – particularly after introducing a random block playback design and nose poke readout to sustain and gauge motivation, respectively.

One further innovation allowed us to circumvent a common limitation in rodent USV studies: emissions are difficult to localize to specific animals.^42^ We made sure that our playback used recordings from males who were stimulated with the *same* female, so that any female-emitted vocalizations would be acoustically similar on both speakers and thus should not be the basis for a preferential response.

Furthermore, playbacks were counterbalanced so that if one female heard audio files from her partner and a stranger male, then that stranger’s partner heard the same two files in her own playback session. With this, we still found a consistent nose poke preference towards each female’s *own* partner, making it unlikely that the stimulus female’s vocalizations drove behavioral discrimination. We therefore conclude that female prairie voles can recognize their partners’ vocalizations, establishing a behavioral foundation for uncovering the neural mechanism of individual vocal recognition.

Rodents are already widely studied to elucidate neural underpinnings of social behavior and recognition. For instance, “nepotopy” – wherein cells responsive to non-kin versus kin-related stimuli are spatially organized – was discovered in the rat lateral septum.^43^ Neurons with causal roles in social recognition memory were found in mice by manipulating circuits between the hippocampus and the lateral septum or nucleus accumbens.^44–46^ Furthermore, neuromodulatory mechanisms to establish long-term preferences for specific individuals were uncovered by studies in prairie voles, which revealed the importance of the oxytocin and vasopressin neuropeptide systems^47–49^ and how they change with experience.^20,50^ Despite this broad neurobiological foundation, no work has explored the neurobiological basis for individual vocal recognition in rodents because the behavioral evidence for it has been lacking. This is likely because vocal differences are instead typically attributed to species, strain, sex, arousal, or social context – not individual identity.^51,52^ Our paradigm in prairie voles creates an opportunity to find neural correlates of individual voices, as has been found in primates,^53,54^ and track their integration into the social recognition circuitry that drives behavioral responses.^55^ Intriguingly, prairie voles have an unusually large auditory cortex compared to other rodent species,^56^ which could reflect an evolutionary adaptation to enhance acoustic cues in the vocalizations from one’s monogamous partner.

Finally, elucidating the basic science of individual vocal recognition offers translational potential. A relatively understudied clinical deficit known as phonagnosia manifests as an impairment in recognizing people by their voice.^57^ Patient studies suggest a neural origin starting within the human temporal lobe and extending beyond it depending on whether there are dysfunctions in processing basic vocal features or the sense of familiarity generated by a vocal percept.^58^ While species differences would need to be factored in, establishing an ethologically grounded behavioral paradigm for individual vocal recognition in rodents will make future studies of the underlying voice perception-to-action circuits possible.

## MATERIALS AND METHODS

### Procedures

Experiments were conducted in strict accordance with the guidelines established by the National Institutes of Health and approved by Emory University’s Institutional Animal Care and Use Committee.

#### Subjects

Adult prairie voles (P60+) were used to assess the role of ultrasonic vocalizations (USVs) as a means of partner identification. 17 voles were used for behavioral experiments. All animals originated from a laboratory breeding colony derived from field-captured voles in Champaign, Illinois. Animals were housed with a 14/10 h light/dark cycle at 68-72°F with *ad libitum* access to water and food (Laboratory Rabbit Diet HF # 5326, LabDiet, St. Louis, MO, USA). Cages contained Bedo’cobbs Laboratory Animal Bedding (The Andersons; Maumee, Ohio) and environmental enrichment, which included cotton pieces to facilitate nest building. Animals were weaned at 20-23 days of age then group housed (2-3 per cage) with age- and sex-matched pups. Experiments occurred during the light cycle (between 9 a.m. and 5 p.m.).

All females were ovariectomized prior to experiments. Females were then primed with subcutaneous administration of estradiol benzoate (17-β-Estradiol-3-Benzoate, Fisher Scientific, 2 μg dissolved in sesame oil) for the 3 days preceding any days on which they were recorded.

#### Data Collection

All recordings were conducted in a designated behavioral-recording room separate from the animal colony. To record socially-induced USVs, males were first removed from their home cage and placed into a plexiglass recording chamber (24.5 × 20.3 mm) lined with clean Alpha-DRI bedding. Males were recorded in three social settings. First on Day 0, a stimulus female (the same stimulus female was used with all males) was placed into the arena and free interaction with the male was audio and video recorded for 30 minutes. The following day, Day 1, males were recorded for 30 minutes with the female that he would be pair-housed with. The male and female were then removed from the recording arena and placed into a shared home-cage, where they remained for 7 days. On Day 8, the male and female were separated. On Day 9, the female was brought back for a playback experiment (outlined below). Later on Day 9, the pair-housed male and female were reintroduced in the recording arena and audio and video data was recorded for another 30 minutes. During all recordings, animals had *ad lib* access to water gel (Clear H2O Scarborough; Scarborough, ME) and food (Laboratory Rabbit Diet HF # 5326).

A microphone (Avisoft CM16/CMPA microphone) was placed above the chamber to record audio data. Audio data were sampled at 300 kHz (Avisoft-Bioacoustics; Glienicke, Germany; CM15/CMPA40-5V), and an UltraSoundGate (116H; Avisoft-Bioacoustics; Glienicke, Germany) data acquisition system was used and integrated with Avisoft-RECORDER software to store the data. A video camera (Canon Vixia HF R800) recorded a top-down view of the chamber at 30 frames-per-second.

#### Vocal Extraction

To extract vocalizations and vocal segments (continuous units of sound), audio files were processed with USVSEG, an open-source MATLAB-based program for detecting and extracting USVs.^59^

Files were bandpass filtered between 15 and 125 kHz, then characterized with a timestep of 0.5 ms. Sounds with fewer than 6 samples (i.e., shorter than 3 ms) were excluded. USVSEG was modified as reported previously^41^ to generate frequency contours – traces of the time, sound frequency, and sound amplitude at each sample within all extracted vocalizations. The contours were then further refined using custom-written MATLAB scripts.

To delineate time blocks where vocalizations occurred in a recording, we ran files through USVSEG and a postprocessing script to generate a structure indicating the time of each USV emission. The files were then broken up into two second intervals and labelled as ‘background noise’ (no vocalizations present) or ‘USV-containing’ (one or more USVs present). These two second intervals were used to generate our playback files below.

#### Vocal Playback

To characterize female interest in USVs, females were placed into a plexiglass arena (20 cm × 24.6 cm) which had a speaker behind an opaque barrier on the left and right sides. The barrier contained holes 1 cm in diameter which the females could poke their nose through.

For each playback session, females were acclimated in the arena while 10 minutes of background noise (see below) played. The acclimation file was generated by finding all two-second intervals without vocal emissions in a control audio file (a male-female interaction not used for our experiment) and combining a random subset of the intervals into a 10-minute acclimation file solely containing background noise. This file was then played back on both speakers simultaneously during the acclimation period prior to each recording.

We used audio data from the male-stimulus pairings on Day 0 to generate playback files for preference testing. Day 0 was chosen because all males were interacting with the same stimulus female on Day 0, and thus any contamination from female vocalizations would be from the same female across all playback files.

During the test period, females heard 10 minute audio files, consisting of blocks of background noise and blocks containing USVs. To generate blocks of background noise, 15 unique ‘background-noise’ intervals were randomly selected and combined to generate one 30-second file. This process was repeated 10 times to generate 10 unique background-noise blocks for a single recording. The same process was followed to generate blocks of ‘USV-containing’ files, using two second intervals with vocalizations. These 20 blocks were then consolidated into a single 10-minute playback file using the ordering seen in **Figure 4**. Whether Order A corresponded to the partner file or the stranger file was randomly assigned. Audio files were manually checked by an experienced observer to ensure accurate file generation.

Which speaker the stranger and partner sounds came from was randomized, but sides were kept consistent within an individual playback session to mimic a more realistic scenario wherein the males were not inexplicably teleporting across the arena. Playbacks were counterbalanced such that when possible, two different females heard the same two playback files. E.g., the playback files for the female from pair 1 were from male 1 (partner) and male 3 (stranger). The same files were played back to the female from pair 3, such that the sounds from male 1 became the stranger sounds, and the sounds from male 3 became the partner sounds. One planned pairing did not occur (pair 8), because the male needed to be removed from the study after the Day 0 recordings. Thus, the female from pair 6 heard playback sounds from partner male 6 and stranger male 8, which was not counterbalanced with an eighth partnered female.

#### Behavior Scoring

For each playback recording, an observer used the video data to score each time a female touched either of the two walls containing a speaker with a paw, as well as all times the female nose-poked towards the speaker. The time each behavior began and ended was recorded through Boris behavioral annotation software.^60^ Unfortunately, one video file was corrupted, so the video from pair 7 was not scored. Observers were blind to vocalizer identity as well as when and which speakers were playing vocalizations during scoring. Matlab scripts were subsequently used to align the recorded behaviors to vocal playback on each speaker.

### Analyses

#### Quantifying Acoustic Features of Vocalizations

Using custom written code (MATLAB), the audio data corresponding to each individual vocal segment was used to extract a fundamental frequency, which was fit to a linearly and sinusoidally modulated function (sinFM) with six features:^31^ onset frequency (f_0_), amplitude of frequency modulation (A_fm_), frequency of frequency modulation (F_fm_), sine phase at sound onset (φ), linear rate of frequency change (slope), and length of sound (duration) (**Figure 1B**):

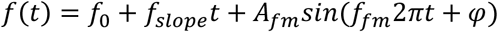

As a way to examine individual variability in vocal features, we compared the distributions of duration and onset frequency across all individual recordings on Day 1 using pairwise Kolmogrov-Smirnov (KS) tests and a posthoc Bonferroni correction for multiple comparisons (uncorrected p < 0.0024 for significance).

#### Characterizing Vocal Space

To visually depict the vocal space across all 6 features of each vocal segment, a t-SNE method of dimensionality reduction was used to project the 6-dimensional vocal representations into a 2-dimensional space (**Figure 2**). This analysis included data from male recordings with his partner (or to-be partner) on Days 9 and 1, respectively. Maps (1001×1001 bins, gaussian smoothed with standard deviation of 34 bins) based on the sinFM parameters were generated^61^ for all males combined (**Figure2A**), all males combined within each social context (**Figure 2B, bottom**), and all males individually within each social context (**Figure 2B, top**). Differences between any two vocal maps were measured by the Jensen-Shannon divergence.^62^

#### Characterizing Acoustic Discriminability

To determine whether individual animal identities could be determined based solely on the features of emitted USVs, we generated a series of 1000 multi-class single vector machine (mcSVM) classifiers. Classifiers were provided with the six sinFM features of each vocal segment plus the identity of the recording pair and tasked with identifying the recording pair given the features of novel vocal segments. For each of the vocal features listed above, values were normalized between 0 and 1 to put all features on the same scale. Then, to train each classifier, 93 vocalizations from each male were randomly selected as training data, and 31 vocalizations were selected as testing data (75 and 25%, respectively, of the least vocal recording, a stimulus recording containing 125 USVs). Each classifier was trained with data from all 7 males, then tested on the ability to predict the identity of the emitter of individual, untrained vocalizations. This was replicated 1000 times per comparison type to generate a range of classifier accuracies.

#### Characterizing Female Behavior

Female behavior during playback was manually scored by a trained observer (see above). Proportions of time responding to USVs as they were playing out from each speaker was then calculated as

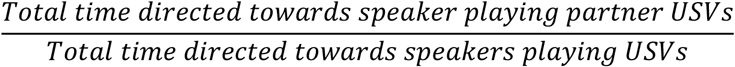

To characterize changes in female interest over time, the total duration of nose-poking was calculated in each 30-second bin. A repeated-measures ANOVA was used to determine an effect of time of nose-poke duration.

### Statistics

Data are represented as mean ± standard deviation unless otherwise indicated. Comparisons across feature distributions were conducted with a one-way ANOVA with a posthoc Bonferroni correction for multiple comparisons. Comparisons between vocal maps were conducted with Jensen-Shannon divergence. Distributions of within-animal divergences in vocal maps to between-animal divergences in vocal maps were compared via a KS test. The distribution of between-animal divergences minus within-animal divergences was compared to a central value of 0 using a ranksum test. Comparisons between female behavioral preferences and chance preferences (50%) used a t-test. The effect of time on nose-poking behavior was assessed with a repeated-measures ANOVA. A p-value of 0.05 was used as the threshold for significance. All statistical analyses were conducted in MATLAB (Mathworks). Data are represented as mean±standard deviation unless otherwise indicated.

## ACKNOWLEDGEMENTS

We thank Lorra Julian and the Emory Primate Center veterinary and animal care staff for vole care. We thank Danial Arslan and Jim Kwon for setting up the ultrasound recording and playback system.

